# A straightforward cell culture insert model to incorporate biochemical and biophysical stromal properties into transplacental transport studies

**DOI:** 10.1101/2024.04.19.590317

**Authors:** Katherine M. Nelson, Bryan J. Ferrick, Hassan Karimi, Christine L. Hatem, Jason P. Gleghorn

## Abstract

**Introduction:** The placental extracellular matrix (ECM) dynamically remodels over pregnancy and in disease. How these changes impact placental barrier function is poorly understood as there are limited *in vitro* models of the placenta with a modifiable stromal compartment to mechanistically investigate these extracellular factors. We developed a straightforward method to incorporate uniform hydrogels into standard cell culture inserts for transplacental transport studies.

**Methods:** Uniform polyacrylamide (PAA) gels were polymerized within cell culture inserts by (re)using the insert packaging to create a closed, controllable environmental chamber. PAA pre-polymer solution was added dropwise via a syringe to the cell culture insert and the atmosphere was purged with an inert gas. Transport and cell culture studies were conducted to validate the model.

**Results:** We successfully incorporated and ECM functionalized uniform PAA gels to cell culture inserts enable cell adhesion and monolayer formation. Imaging and analyte transport studies validated gel formation and expected mass transport results and successful cell studies confirmed cell viability, monolayer formation, and that the model could be used transplacental transport studies. Detailed methods and validation protocols are included.

**Discussion:** It is well appreciated that ECM biophysical and biochemical properties impact cell phenotype and cell signaling in many tissues including the placenta. The incorporation of a PAA gel within a cell culture insert enables independent study of placental ECM biophysical and biochemical properties in the context of transplacental transport. These straightforward and low-cost methods to build three dimensional cellular models are readily adoptable by the wider scientific community.

## Introduction

Mechanistic understanding of transplacental transport is essential for a variety of homeostatic and disease processes, including nutrient and waste transport, drug delivery, inflammatory response, and infection. A commonly used method to study these processes is with a cell culture insert (e.g., Transwell insert), which uses a polyethylene terephthalate (PET) or polycarbonate membrane to support monolayer cell culture and generate separate reservoirs of apical and basal cell culture media within a multi-well cell culture plate (1– 7). While these inserts provide a simple, reliable, and easily accessible system to study transplacental transport, they do not incorporate physiologically relevant stromal properties known to affect tissue barrier properties and cell function. It is well recognized that the extracellular matrix (ECM) is dynamic, and the biophysical properties (e.g., stiffness, moduli, geometry, etc.) and biochemical properties (e.g., protein composition, ECM organization, etc.) of the placental ECM change over gestation and in disease(8–12). Inclusion of these parameters is limited primarily to three-dimensional cell culture models or “organ-on-a-chip” models (13–17), which significantly increase the complexity and decrease the throughput of the number of individual samples that can be tested simultaneously. To date, independent control of the biophysical and biochemical properties of tissue stroma has not been introduced to cell culture insert systems in a straight-forward and reproducible manner that would make them suitable for transport studies.

Polyacrylamide (PAA) hydrogels have been widely used to allow for independent investigation of the effects of elastic modulus and protein identity on cell function in many systems (18–20). PAA gels are bioinert, and their elastic modulus is a function of the weight percent of polymers added to the pre-gel solution. The polymerized gel surface can be functionalized covalently with selected ECM proteins for cell adhesion and culture on the hydrogel surface as a monolayer or individual cells. PAA gels are classically made by sandwiching the pre-polymer solution between two glass slides or coverslips to form reproducible shapes and heights that are covalently bonded to a single glass coverslip (21). Importantly, this technique also reduces the surface area of the solution exposed to the air, as oxygen is an inhibitor to the gel polymerization reaction and will prevent complete “gelation” of the pre-polymer solution (20). Numerous other hydrogels, synthetic (e.g., polyethylene glycol [PEG], poly(lactic-co-glycolic acid) [PLGA], etc.) and naturally derived (e.g., collagen, hyaluronic acid, alginate, fibrin, etc.), have been used for studies that mimic the ECM and can also allow for embedding stromal cells within the bulk of the hydrogel (22– 25); however, independent control of biophysical properties and protein or peptide identity are difficult and require more care.

To create a successful transplacental transport model with a tunable hydrogel stromal compartment, the model must contain separable apical and basal compartments, a permeable interface between the two, and a hydrogel stromal compartment of uniform and repeatable thickness between individual models. Here, we report a simple model of the placenta with independent control over the ECM biophysical and compositional properties in a cell culture insert for transplacental transport studies (**Figure 1**). We created functionalized PAA hydrogels within the cell culture insert as an example stromal compartment and provided validation data and detailed protocols (**Supplemental Material**) for users to make this model with corresponding experimental checks to validate successful fabrication and performance in their laboratory. Given the insert geometry and the permeable membrane, traditional methods to create these PAA gels cannot be adapted to form a PAA hydrogel within a cell culture insert. As such, we developed an inexpensive, simple method to create uniform thickness hydrogels *in situ* within cell culture inserts using dropwise addition of prepolymer and polymerization within the purged atmosphere of the cell culture insert packaging. Notably, the methods used to create a reproducible PAA gel of uniform thickness can be extended to other types of hydrogels to enable the straightforward investigation of additional effects on barrier function, including cell-cell interactions and reciprocal signaling between cell populations with the incorporation of additional cell types.

**Fig. 1.**
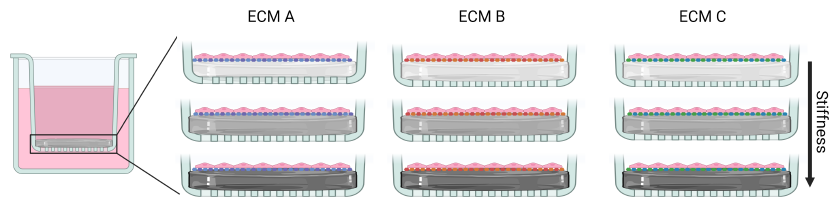
ECM stiffness and composition can be independently tuned in a cell culture insert model to study their effects on transport.

## Materials and Methods

### In situ fabrication of a PAA gel in a cell culture insert

Acrylamide or N-hydroxysuccinimide (NHS)-acrylamide, bis-acrylamide, ammonium persulfate (APS), and Tetramethylethylenediamine (TEMED) were mixed following standard methods to create a PAA gel **Sup. Protocol 1** (21, 26). The solution was aspirated into a 1 mL syringe with a 23G needle. The cell culture insert (Cell Treat or Corning Falcon) was kept within the sterile packaging to create a semi-closed environment to ensure proper polymerization. The packaging was sprayed with 70% ethanol and allowed to air dry. The removable paper seal of the packaging was punctured with the needle attached to the syringe with the pre-polymer solution. For a 12-well cell culture insert, we quickly added 100 μL of pre-polymer solution dropwise to the insert, removed the prepolymer syringe from the package, and introduced a needle attached to an inert gas (nitrogen or argon) source by puncturing a separate hole in the paper packaging. The atmosphere within the packaging was purged with inert gas to displace the air (oxygen) for 90-120 sec. The inserts were left undisturbed upright for 30 minutes to fully polymerize the gels before being moved into a biosafety cabinet and removed from the packaging. At this point, the cell culture inserts underwent the ECM functionalization procedure or were stored at 4ºC with the hydrogel fully wetted by PBS in the basal and apical compartments.

### PAA ECM functionalization

There are two standard methods used to functionalize PAA gels: (1) addition of a heterobifunctional linker (sulfo-SANPAH) with one end that can be covalently coupled to the PAA gel surface via UV and a reactive N-hydroxysuccinimide (NHS) group on the other end which can freely react with primary amines in proteins (21) or (2) incorporation of a reactive NHS group into the gel using an NHS-acrylamide (NHS-AA) monomer (26). Both strategies result in PAA hydrogels with functional reactive NHS groups that covalently attach proteins to the gel surface. For the sulfo-SANPAH functionalization, gels were washed, and sulfo-SANPAH was added to the gel surface and exposed to UV light. Next, excess sulfo-SANPAH was washed away, and 300 μL of 150 μg/mL type I rat tail collagen was added to the gel surface, incubated overnight at 4ºC, and washed with PBS. For the NHS-AA method, a portion of the acrylamide was replaces with NHS-AA (6:1 ratio of acrylamide to NHS-AA) creating the gels. Type I rat tail collagen (300 μL of 150 μg/mL) was added to the surface and incubated overnight at 4ºC with a subsequent PBS wash for covalent attachment of the collagen-I to the hydrogel surface.

### In situ imaging of PAA hydrogels within cell culture inserts

To image the gel thickness, 1 μm fluorescent beads (Bangs Laboratories) were added to the pre-polymer solution and polymerized into the gel’s bulk to enable visualization via fluorescence microscopy. The gel-laden insert was placed on a coverslip, and a z-stack of images (Zeiss, LSM 800 confocal microscope) was taken in multiple regions of interest over the insert to ensure the uniformity of the gel.

### Cell culture and immunofluorescence staining

BeWo b30 cells were maintained in F-12K supplemented with 10% FBS and 1% penicillin/streptomycin at 37ºC. After ECM functionalization, BeWo b30 cells were seeded onto the surface of the gel at 2x10^5^ cells/cm^2^ in complete media. Cells were left to adhere to the PAA gel surface for 4 hours before replacing the apical cell culture medium. The medium in the apical and basal compartments was changed daily over culture for 4-6 days post-seeding. Following culture, cells were fixed with 4% paraformaldehyde (PFA), permeabilized with 0.1% triton-x, and stained with DAPI, phalloidin 647 (Abcamab176759), rabbit anti-E-cadherin (Cell Signaling-3195S) and donkey anti-rabbit 488 (ImmunoReagents-F488NHSX). The insert membrane was cut with a scalpel to disconnect it from the insert walls so the membrane-gel unit could be removed and placed cell side down on a glass coverslip to enable imaging via Zeiss LSM 800 confocal microscope.

### Passive diffusion assay for barrier function assessment

Medium in the basal well was replaced with fresh cell culture medium and medium in the apical well was replaced with medium supplemented with either 200 μg/mL FITC or 1.5 mg/mL 150 kDa FITC-dextran. Complete protocols for this assay can be found in **Sup. Protocol 2**. At designated time points after adding fluorescent molecules, 50 μL of the basal solution was sampled in triplicate and replaced with fresh cell culture medium. The fluorescent intensity of the samples was measured on a plate reader, and concentration was calculated from a standard curve and normalized to the initial known apical concentration.

## Results

### Fabrication of a uniform PAA gel in a cell culture insert

Given that PAA gel polymerization needs to occur in the absence of oxygen, we sought to identify a simple procedure to control the atmosphere around the cell culture insert to enable *in situ* polymerization of the gel on the cell culture insert membrane. Individually packaged cell culture inserts come in a molded plastic case with a backing paper on one side that is peeled off to remove the cell culture insert. We chose to keep the packaging intact and gain access to the cell culture insert by puncturing through the paper backing to create a sterile and enclosed environment to create our gels (**Figure 2AB, Sup. Protocol 1**). The polymer solution was carefully but quickly added dropwise to the insert through a syringe, puncturing the paper seal. After addition, the syringe was removed, and the resulting hole was used to insert a needle attached to an inert gas (nitrogen or argon). The packaging was purged with the gas to displace atmospheric oxygen for 1-2 minutes, and the packaging was undisturbed to allow for PAA gel polymerization. After 30 minutes, the gel was fully polymerized, and the insert was removed from the packaging to store fully hydrated in sterile PBS or immediately prepared for covalent chemical conjugation of proteins of interest to the gel surface. It is important to note that the PAA prepolymer solution will immediately start to polymerize once the initiators (APS and TEMED) are added; therefore, after mixing, it is essential to work quickly from loading the syringe to dropwise addition into the cell culture inserts.

**Fig. 2.**
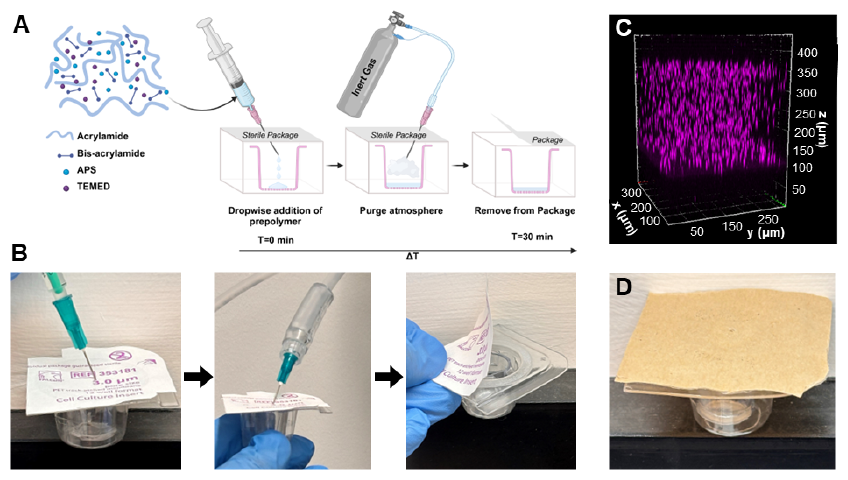
Uniform PAA gels were successfully created *in situ* in a cell culture insert. (A) Schematic of the method developed and (B) corresponding photographs of experimental method. (C) 3D reconstructed view of fluorescent microspheres embedded throughout the bulk of a uniform PAA gel within a cell culture insert. (D) picture of re-used packaging with a new paper back secured with double-sided tape.

To visualize the polymerized gel, we added 1 μm fluorescent microspheres into the pre-polymer solution to be distributed throughout the bulk of the gel. A three-dimensional reconstruction of z-stacked fluorescent images of the PAA gel within the cell culture insert revealed a uniform, flat hydrogel was created over the entire cell culture area (**Figure 2C**). This process was repeatable over multiple cell culture inserts, generating flat PAA gels with uniform thickness and dimensionality. We further tested different PAA hydrogel formulations (**Table 1**) that would match a range of placental ECM stiffnesses found in a healthy or dysfunctional placenta (1 kPa, 10 kPa, and 20 kPa) (21). We also tested the primary formulation (using acrylamide monomer (21)) and an alternative formulation (use of an NHS-coupled acrylamide monomer (26)) for PAA gels, which use different functional chemistry to covalently couple proteins to the gel surface (additional details in methods: PAA ECM functionalization). The gel fabrication method developed herein was compatible with all the PAA gel formulations tested and produced uniform replicable PAA gels with defined thicknesses across multiple cell culture inserts.

**Table 1.**
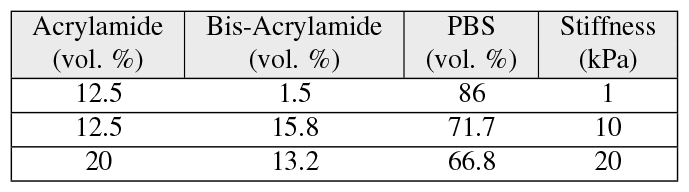
Formulations by volume percent of acrylamide, bis-acrylamide, and PBS used to make PAA gels of 1, 10, and 20 kPa stiffnesses.

| Acrylamide (vol. %) | Bis-Acrylamide (vol. %) | PBS (vol. %) | Stiffness (kPa) |
| --- | --- | --- | --- |
| 12.5 | 1.5 | 86 | 1 |
| 12.5 | 15.8 | 71.7 | 10 |
| 20 | 13.2 | 66.8 | 20 |

### Generalizability of the methods

In our search for commercial cell insert vendors, we found that products were packaged either individually or with multiple in a multi-well plate format packaged together in a bulk format. To determine if our method could be easily adapted to these types of inserts at a low cost, we attempted to reuse individual packaging for PAA gel polymerization in new cell culture inserts (**Figure 2D**). A multi-cell culture insert package was opened in the biosafety cabinet, and standard ethanol sterilization and aseptic techniques were used to transfer the cell culture insert to a previously opened individual package. Double-sided tape was placed around the rim of the packaging, and a new piece of paper was affixed to the packaging to replace the original paper backing. Once assembled, the same methods successfully created PAA gels *in situ*.

### Validation and characteristics of a gel-cell culture insert system as a transport model

Traditionally, in transcellular transport models using cell culture inserts, transepithelial electrical resistance (TEER) or fluorescent molecules with well-defined permissivity or resistance to transport are used to assess the established cell monolayer’s initial validity and barrier function. Indeed, these same methods are compatible with cell culture inserts that contain a hydrogel. However, the diffusivity and diffusion times of fluorescent species from the apical to basal compartment will be a function of the combined contributions of the cell monolayer barrier function and the physical properties of the hydrogel, including the gel thickness, porosity, and pore size, and the concentration gradient produced. To provide helpful user benchmarks, we focused on characterizing apical to basal transport of two fluorescent molecules of varying size with and without a monolayer to characterize relevant timescales and contributions of the PAA gel and trophoblast monolayer in this model. These data and protocols **Sup. Protocol 2** can be used to validate correct PAA gel fabrication and establishment of the model to subsequently test the performance of known species that should not cross a functional trophoblast layer (e.g., insulin, heparin, etc.), can readily pass (e.g., sodium fluorescein, antipyrine, etc.), and those that are actively transported (e.g., glucose, amino acids, etc.).

To validate that the polymerized gel was properly formed and sealed around the walls of the cell culture insert, fluorescent tracer diffusion of small (FITC) and large (150 kDa FITC-dextran) was quantified and compared between inserts with PAA gels and inserts without a gel as a control. As expected, in non-PAA gel controls, FITC readily diffused across the membrane and could be measured in the basal compartment within 10 min of addition (**Figure 3A**). FITC-dextran, a much larger molecule, similarly diffused into the basal compartment quickly in non-gel controls (**Figure 3B**). Adding the PAA gel significantly increased the diffusion times of fluorescent molecules. The diffusion assay needed to be performed for 2.5 hours before a meaningful signal was measured in the compartment for FITC and 24 hours for FITC-dextran. A similar and expected size-dependent difference in transport rates between FITC and FITC-dextran were observed as in controls, with the diffusion times between the two species increasing dramatically due to the presence and properties of the hydrogel in the cell culture insert. These data confirm the expected effects on the transport properties due to the PAA gel and similarly confirm that the hydrogel is formed such that it has an intimate connection to the cell culture insert walls and membrane, requiring transport through the gel with no opportunity to flow around the gel to the basal compartment.

**Fig. 3.**
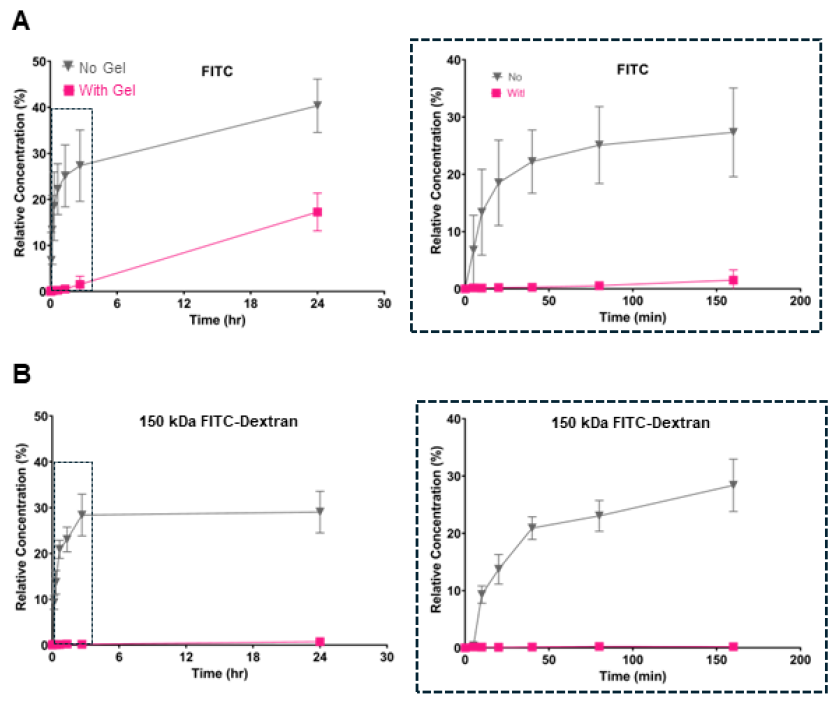
Quantitation of fluorescent analyte transport demonstrate successful PAA gel fabrication and seal to the insert wall. Basal compartment concentration relative to starting apical compartment concentration of (A) FITC or (B) 150 kDa FITC-dextran in PAA gel inserts and non-PAA gel insert controls over 24 hrs. Corresponding inset view of plot on right in dashed box. n=3. Error bars represent standard deviation.

### Validation of BeWo monolayer formation and barrier function

PAA gels are inert and must be coated with ECM proteins, individually purified or complex mixtures, to enable cell adhesion and growth. This is done with a covalent coupling scheme using a heterobifunctional linker or an NHS-coupled acrylamide. To ensure standard functionalization protocols and different PAA gel formulations were compatible with cell culture in these cell culture inserts, type I rat tail collagen was covalently bonded to the surface of the PAA gels. BeWo b30 cells were subsequently seeded onto functionalized PAA gels in cell culture inserts. To visually confirm monolayer formation following 4-6 days of culture, the cells were fixed and immunofluorescently stained for nuclear (DAPI) and junctional (E-cadherin) markers (**Figure 4A**). A complete monolayer was confirmed by densely packed cells and junctional protein staining between cells in numerous fields of view across the entire PAA gel surface. As these studies aimed to confirm viable cell culture, monolayer formation, and barrier function of a widely used trophoblast cell line, forskolin was not added to the cell culture medium to induce fusion but could be for functional studies. The barrier function of the BeWo b30 cell monolayers was functionally validated using a 150 kDa-FITC-dextran diffusion assay (**Figure 4B, Sup. Protocol 2**). Diffusion of FITC-dextran across a cellularized gel-laden insert was compared to acellular non-PAA gel insert control. Inserts with cells had no detectable FITC-dextran transport after 72 hours, while inserts without gels had reached equilibrium by 24 hours. This indicates that BeWo b30 cells create a functional monolayer on collagen-I coated PAA gels and that ECM functionalization of the PAA gels in a cell culture insert was successful.

**Fig. 4.**
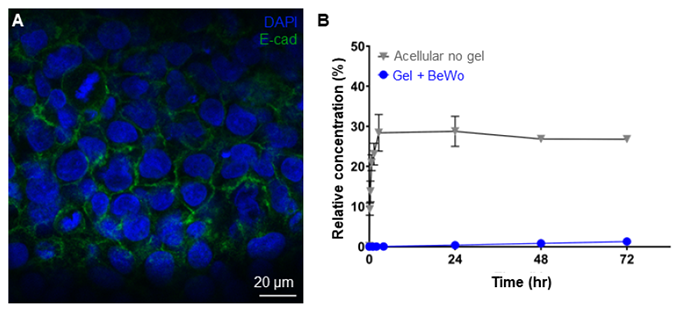
BeWo b30 cells generate a functional monolayer on PAA gel inserts. (A) Immunostained image of a BeWo b30 monolayer before fusion on a 20 kPa PAA gel in an insert. (B) Basal compartment concentration relative to the starting apical compartment concentration of 150 kDa FITC-Dextran across a BeWo b30 monolayer on a 20 kPa PAA gel in an insert over 72 hrs of culture. n=3 for all except n=1 for no gel 48 and 72 hrs. Error bars represent standard deviation.

## Discussion

The placental ECM biophysical and biochemical properties dynamically change throughout gestation and in disease, yet how these factors and the mechanisms they regulate influence transplacental transport remains unknown due to limited models. ECM proteins have been shown to be altered in diseases of the placenta, which may contribute to placental dysfunction. Collagen-I and laminin β1 and β2 are upregulated in preeclampsia, while collagen-IV and fibronectin are downregulated compared to placentas from healthy pregnancies. Interestingly, BeWos cultured with forskolin to induce fusion on either collagen-I or collagen-IV substratum revealed slower fusion rates on collagen-I compared to collagen-IV, indicating that specific ECM proteins may play a role in pathology etiology (27). Similarly, different placental stiffness is found in disease; using Doppler ultrasound, healthy placental stiffness was measured at 2-10 kPa while preeclamptic placentas average 20-25 kPa (8, 10). Importantly, these mechanical stiffness ranges cannot be achieved with numerous hydrogel formulations, several of which, like type I collagen gels, are an order of magnitude softer than healthy placenta (28). However, PAA gels can be reliably created from 0.2-300 kPa by altering the ratio of acrylamide to bis-acrylamide to PBS of the gel solution (20, 21, 29). Mechanistically, PAA hydrogels have been used to study the ability of trophoblasts to fuse on different stiffnesses. On PAA gels coated with collagen-I, increased elastic modulus (17 kPa) decreased the ability of BeWos to fuse and form a syncytium and decreased the amount of human chorionic gonadotropin (hCG) expressed by the cells compared to cells cultured on softer, 7 or 1.3 kPa, substrates, respectively (30). These data highlight the need for functional models that can be made simply and at a low cost that incorporate and dissect the relative contributions of the ECM to placental transport.

We developed a simple method to create uniform, reproducible hydrogels within a cell culture insert. Most cell culture insert assays use a protein coating on the membrane as the cell substratum (1, 3, 4) or, infrequently, an ECM gel without regard for stiffness (2, 7). Neither approach is amenable to studying the extracellular environment’s biophysical effects, including stiffness, on placental barrier function. By keeping the insert within the sterile packing, purging the system with inert gas (nitrogen or argon), and adding the polymer solution dropwise via a syringe and needle, a uniform PAA gel can be created in an insert (**Figure 2**). Whereas both nitrogen and argon will work if the purge is long enough (minutes), argon is denser than air, so it will displace the air faster and more reliably than nitrogen. However, nitrogen is typically less expensive and more readily available, and we confirmed it can be successfully used. Using published formulations, these methods can create PAA gels ranging from at least 1-200 kPa. The thickness of the PAA gel is a function of the volume added and the cell culture insert’s surface area. However, the smaller the volume used, the larger the meniscus and curvature of the gel. We found that 90 μl/cm^2^ cell culture insert provided thick enough gels to ensure the elastic modulus experienced by the cells was due to the gel only (31, 32), but a large enough volume to ensure reproducibly uniform thickness gels. Lastly, we also validated that these methods are extensible to any closed environmental chamber that allows dropwise addition of hydrogel pre-polymer and purging of atmospheric oxygen to create uniform, fully polymerized hydrogels *in situ* within an insert. The straightforward fluorophore diffusion assays outlined here can easily be completed in any system before an experiment starts to confirm the integrity of the gel-insert interface and the cell monolayer. We focused on defining assays geared toward the end user to confirm gel fabrication and establish the culture model in an individual lab. As such, we show the results of two different-sized fluorescent analytes that can be used to validate gel fabrication and complete sealing of the gel to the insert sidewalls (**Figure 3**), as well as the performance of an unfused BeWo b30 monolayer that can be used to validate successful surface protein conjugation and tissue barrier function (**Figure 4**). Absolute analyte concentrations in the basal compartment can be normalized to the starting apical concentration to compare across different experimental groups. For quantitative measures of diffusivity, however, the effects of dilution from temporal sampling need to be accounted for. It is also important to note that the pore size of the PAA gel will change with the starting monomer concentration and, therefore, is related to the gel stiffness (33). This will affect analytes’ transport through gels of different stiffnesses, resulting in different apparent rates. Therefore, the results of time-dependent studies should account for these differences. In addition to analyte diffusion, transepithelial electrical resistance (TEER) is compatible with these gel-laden inserts as commercial instruments are built and calibrated for Transwell inserts. However, when inserting the electrode into the insert, caution must be taken to ensure the bottom of the electrode does not touch the surface of the gel. This can be accounted for by simply adjusting the probes’ length before use.

Whereas we demonstrate a simple monolayer of BeWo cells on the surface of a collagen-I coated PAA gel, standard methods to add additional cell types (34) and proteins to cell culture insert models can also be applied to this system. Other proteins, such as laminin, fibronectin, collagen-IV, or decellularized placental ECM, can be covalently linked to the PAA gels with this chemistry to elucidate their specific roles. Various cells can be added, including primary trophoblasts co-cultured with endothelial cells or fibroblasts on the underside surface of the insert, to investigate intercellular signaling and relative contributions to placental barrier function. In addition, synthetic hydrogel systems with tunable stiffness that have been designed to sustain viable cells after encapsulation can be produced in an insert using these methods, given the added control over the atmosphere during polymerization. These types of gels enable the addition of encapsulated cells within the bulk, which could be leveraged to include stromal cells, such as placental fibroblasts or Hofbauer cells, amongst others, to expand the applicability of this simple 3D culture model to delineate the role of cell-cell and reciprocal signaling in transplacental transport studies. This method can also be used to create uniform gels for natural hydrogels, including collagen-I or Matrigel; however, note that these gels are significantly softer than placental tissue (28) (**Figure 5**).

**Fig. 5.**
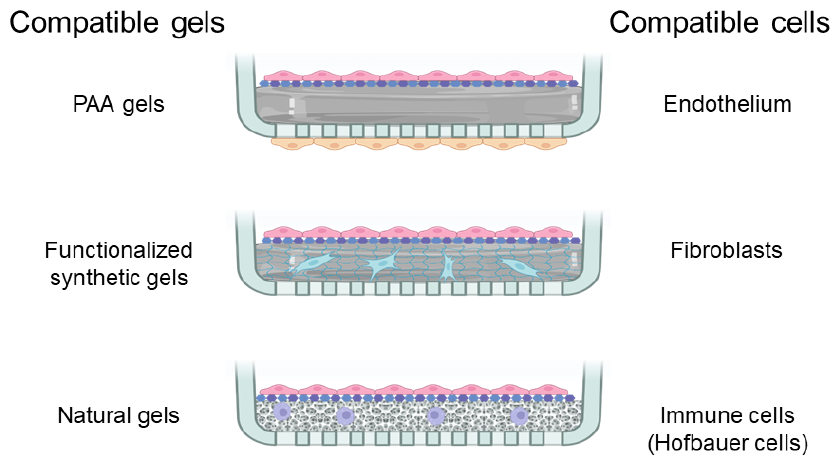
Examples of other types of gel systems, cells, and cell configurations that would be compatible with gel fabrication methods herein.

As we further elucidate the mechanisms behind the placenta’s barrier function, understanding the role of the ECM in this vital function during both homeostasis and stress is necessary. This will allow for novel therapeutics for pregnant people (35, 36) to treat ECM-related conditions and improve treatment with known therapies. The model and techniques we present here are simple and easily adaptable to study a wide variety of questions regarding the mechanisms underpinning transplacental transport.

## ACKNOWLEDGEMENTS

This work was supported in part by grants from the National Institutes of Health: T32GM133395 (KMN), F31HD105398 (KMN), and U19AI158930. Figures were made with BioRender. BioRxiv template adapted from Henriques lab.

## AUTHOR CONTRIBUTIONS

Conceptualization (KMN, JPG), Methodology (KMN, HK, CLM, JPG), Investigation (BJF, HK, CLH), Validation (KMN, JPG), Visualization (KMN, BJF, HK, JPG), Formal Analysis (KMN, BJF, HK, JPG), Writing – Original Draft (KMN), Writing – Review & Editing (BJF, HK, CLM, JPG), Supervision (JPG), Project Administration (KLM, JPG), Funding acquisition (KLM, JPG)

## Supplementary Note 1

### Protocol 1: Polymerization of flat, uniform PAA gel in a cell culture insert

#### Materials needed

PAA prepolymer solution | Cell culture inserts in sterile packaging (Cell Treat, Corning Falcon) | 1 ml syringe | 23G needle | Inert gas (Nitrogen or Argon) | 1.5 ml Centrifuge tube

##### Procedure

1. Prepare the PAA polymer solution for desired stiffness based in a centrifuge tube [17,18].
  - Benchmark values for physiological placental stiffness in Table 1.
2. Quickly aspirate desired volume for number of gels to create into a 1 ml syringe and affix the needle.
  - 90 μl/cm2 or 100 μl/12 well insert
3. Insert the syringe needle into the insert packaging halfway into the insert.
4. Gently add the prepolymer solution onto the insert’s membrane drop by drop.
  - After mixing the polymer solution it will immediately begin to gel, so the solution might gel in the syringe if this is not done quickly.
5. Remove the syringe and insert needle with inert gas (N2 or Argon) line into the packaging to purge the atmospheric oxygen from the system.
  - Make sure the needle is pointed towards to wall of the insert to not disrupt the surface of the pre-polymer/gel.
6. Purge for 90s-120s.
  - The gel should start to polymerize by this point.
7. Remove syringe and let the gel fully polymerize for 30 minutes.

## Supplementary Note 2

### Protocol 2: Diffusion Assay

#### Materials needed

Cell culture insert with PAA gel | 12-well plate | Fluorescent analyte (e.g., 70 or 150 KDa FITC-Dextran) | FITC | Cell culture media | 96-well plate | Plate reader

##### Procedure

1. Make a stock solution of the fluorescent analyte in cell culture media.
  - FITC-Dextran > 0.667 mg/ml
  - FITC > 133.33 μg/ml
2. Make a standard curve of the analyte.
  - Starting at the max concentration, make serial dilutions and pipette 100 μL of each standard in triplicate into a well in the 96-well plate.
  - Add 100 μL in triplicate of the media solution added to the apical chamber to the plate if you wish to normalize basal concentration to starting apical concentration.
  - Store the well plate in the dark.
3. Remove medium (if any) from the basal compartment and add 1 ml of culture media into basal well.
4. Remove medium (if any) from the apical compartment and add 0.5 ml cell culture media with the analyte into apical compartment
  - The goal is to make sure the heights of the fluid in each compartment are the same so there is minimal pressure driven flow. Adjust volumes as needed to achieve this.
  - Due to pressure gradients created by the fluid, it is important to add basal medium while fluid is in the apical compartment to not disrupt the cell layer.
  - To ensure diffusion is accurately measured, replace basal medium before replacing apical medium with analyte.
5. Sample 50 μl of basal solution at various time points (ex: 1, 6, 12, 24, 72, 120, 1440 min) in triplicate. Replace the volume removed with fresh media.
6. Add samples into the 96-well plate to be read on plate reader.
  - Samples will be sensitive to light so store the plate in the dark.
7. Add 50 μl plain cell culture media to each well to bring the volume to 100 μl to get a more accurate reading.
8. Read each 96-well plate with the plate reader set at the correct excitation/emission wavelengths for the analyte you used (e.g., FITC (498/517)).
  - Ensure the samples for the standard curve are on the same plate.
  - Reading can be done after each sample collection with a new standard curve for each timepoint or samples can be stored in the dark and read at the end of the experiment.
  - Photobleaching can occur so ensure each well that needs to be measured is only read one time.
9. Calculate the analyte concentration in each sample to generate a concentration vs time plot that can be analyzed as a validation of model set-up. If dilutions at each point are accounted for at each timepoint, properties including analyte diffusion rates can be calculated.

## References

1. Michael K Wong, Edward W Li, Mohamed Adam, Ponnambalam R Selvaganapathy, and Sandeep Raha. Establishment of an in vitro placental barrier model cultured under physiologically relevant oxygen levels. Molecular Human Reproduction, page gaaa018, March 2020.

2. Navein Arumugasaamy, Alana Gudelsky, Amelia Hurley-Novatny, Peter C. W. Kim, and John P. Fisher. Model Placental Barrier Phenotypic Response to Fluoxetine and Sertraline: A Comparative Study. Advanced Healthcare Materials, 8(18):1900476–1900476, September 2019.

3. Katherine C. Fein, Mariah L. Arral, Julie S. Kim, Alexandra N. Newby, and Kathryn A. Whitehead. Placental drug transport and fetal exposure during pregnancy is determined by drug molecular size, chemistry, and conformation. Journal of controlled release: official journal of the Controlled Release Society, 361:29–39, September 2023.

4. Hequn Li, Bennard van Ravenzwaay, Ivonne M. C. M. Rietjens, and Jochem Louisse. Assessment of an in vitro transport model using BeWo b30 cells to predict placental transfer of compounds. Archives of Toxicology, 87(9):1661–1669, September 2013.

5. Leonie Aengenheister, Kerda Keevend, Carina Muoth, René Schönenberger, Liliane Diener, Peter Wick, and Tina Buerki-Thurnherr. An advanced human in vitro co-culture model for translocation studies across the placental barrier. Scientific Reports, 8(1):5388–5388, December 2018.

6. Marie Sønnegaard Poulsen, Erik Rytting, Tina Mose, and Lisbeth E Knudsen. Modeling placental transport: Correlation of in vitro BeWo cell permeability and ex vivo human placental perfusion. Toxicology in Vitro, 23:1380–1386, 2009.

7. Navein Arumugasaamy, Leila E. Ettehadieh, Che-Ying Kuo, Dominic Paquin-Proulx, Shannon M. Kitchen, Marco Santoro, Jesse K. Placone, Paola P. Silveira, Renato S. Aguiar, Douglas F. Nixon, John P. Fisher, and Peter C. W. Kim. Biomimetic Placenta-Fetus Model Demonstrating Maternal–Fetal Transmission and Fetal Neural Toxicity of Zika Virus. Annals of Biomedical Engineering, 46(12):1963–1974, December 2018.

8. Fahrettin Kilic, Yasemin Kayadibi, Mehmet Aytac Yuksel, Ibrahim Adaletli, Fethi Emre Ustabasioglu, Mahmut Oncul, Riza Madazli, Mehmet Halit Yilmaz, Ismail Mihmanli, and Fatih Kantarci. Shear wave elastography of placenta: in vivo quantitation of placental elasticity in preeclampsia. Diagnostic and Interventional Radiology, 21(3):202–207, May 2015.

9. Miki Mori, Gen Ishikawa, Shan-Shun Luo, Takuya Mishima, Tadashi Goto, John M. Robinson, Shigeki Matsubara, Toshiyuki Takeshita, Hiroaki Kataoka, and Toshihiro Takizawa. The Cytotrophoblast Layer of Human Chorionic Villi Becomes Thinner but Maintains Its Structural Integrity During Gestation1. Biology of Reproduction, 76(1):164–172, January 2007.

10. Michail Spiliopoulos, Che-Ying Kuo, Avinash Eranki, Marni Jacobs, Christopher T. Rossi, Sara N. Iqbal, John P. Fisher, Melissa H. Fries, and Peter C. W. Kim. Characterizing placental stiffness using ultrasound shear-wave elastography in healthy and preeclamptic pregnancies. Archives of Gynecology and Obstetrics, July 2020.

11. Erbil Karaman, Harun Arslan, Orkun Çetin, Hanιm Güler Şahin, Aydin Bora, Alparslan Yavuz, Sadi Elasan, and İbrahim Akbudak. Comparison of placental elasticity in normal and pre-eclamptic pregnant women by acoustic radiation force impulse elastosonography. Journal of Obstetrics and Gynaecology Research, 42(11):1464–1470, 2016.

12. Blakely B O’Connor, Benjamin D Pope, Michael M Peters, Carrie Ris-Stalpers, and Kevin K Parker. The role of extracellular matrix in normal and pathological pregnancy: Future applications of microphysiological systems in reproductive medicine. Experimental Biology and Medicine, 245(13):1163–1174, July 2020.

13. Mario Cabodi, Nak Won Choi, Jason P. Gleghorn, Christopher S. D. Lee, Lawrence J. Bonassar, and Abraham D. Stroock. A Microfluidic Biomaterial. Journal of the American Chemical Society, 127(40):13788–13789, October 2005.

14. Emily M. Chandler, Caroline M. Berglund, Jason S. Lee, William J. Polacheck, Jason P. Gleghorn, Brian J. Kirby, and Claudia Fischbach. Stiffness of photocrosslinked RGD-alginate gels regulates adipose progenitor cell behavior. Biotechnology and Bioengineering, 108(7):1683–1692, 2011.

15. Joshua T. Morgan, Jasmine Shirazi, Erica M. Comber, Christian Eschenburg, and Jason P. Gleghorn. Fabrication of centimeter-scale and geometrically arbitrary vascular networks using in vitro self-assembly. Biomaterials, 189:37–47, January 2019.

16. Catherine S. Millar-Haskell, Allyson M. Dang, and Jason P. Gleghorn. Coupling synthetic biology and programmable materials to construct complex tissue ecosystems. MRS communications, 9(2):421–432, June 2019.

17. Brea Chernokal, Cailin R. Gonyea, and Jason P. Gleghorn. Lung Development in a Dish: Models to Interrogate the Cellular Niche and the Role of Mechanical Forces in Development. In Chelsea M. Magin, editor, Engineering Translational Models of Lung Homeostasis and Disease, Advances in Experimental Medicine and Biology, pages 29–48. Springer International Publishing, Cham, 2023. ISBN 978-3-031-26625-6. doi: 10.1007/978-3-031-26625-6_

18. Robert J. Pelham and Yu-li Wang. Cell locomotion and focal adhesions are regulated by substrateflexibility. Proceedings of the National Academy of Sciences, 94(25):13661–13665, December 1997.

19. Alex P. Rickel, Hanna J. Sanyour, Neil A. Leyda, and Zhongkui Hong. Extracellular Matrix Proteins and Substrate Stiffness Synergistically Regulate Vascular Smooth Muscle Cell Migration and Cortical Cytoskeleton Organization. ACS applied bio materials, 3(4):2360–2369, April 2020.

20. Catherine S. Millar-Haskell and Jason P. Gleghorn. A Large-format Polyacrylamide Gel with Controllable Matrix Mechanics for Mammalian Cell Culture and Conditioned Media Production. Bio-Protocol, 13(17):e4807, September 2023.

21. Justin R. Tse and Adam J. Engler. Preparation of Hydrogel Substrates with Tunable Mechanical Properties. Current Protocols in Cell Biology, 47(1):10.16.1–10.16.16, 2010.

22. Brielle Hayward-Piatkovskyi, Cailin R. Gonyea, Sienna C. Pyle, Krithika Lingappan, and Jason P. Gleghorn. Sex-related external factors influence pulmonary vascular angiogenesis in a sex-dependent manner. American Journal of Physiology-Heart and Circulatory Physiology, 324(1):H26–H32, January 2023.

23. Jasmine Shirazi, Joshua T. Morgan, Erica M. Comber, and Jason P. Gleghorn. Generation and morphological quantification of large scale, three-dimensional, self-assembled vascular networks. MethodsX, 6:1907–1918, January 2019.

24. Christopher S. D. Lee, Jason P. Gleghorn, Nak Won Choi, Mario Cabodi, Abraham D. Stroock, and Lawrence J. Bonassar. Integration of layered chondrocyte-seeded alginate hydrogel scaffolds. Biomaterials, 28(19):2987–2993, July 2007.

25. Nak Won Choi, Mario Cabodi, Brittany Held, Jason P. Gleghorn, Lawrence J. Bonassar, and Abraham D. Stroock. Microfluidic scaffolds for tissue engineering. Nature Materials, 6(11): 908–915, November 2007.

26. Jun Kumai, Satoru Sasagawa, Masanobu Horie, and Yoshihiro Yui. A Novel Method for Polyacrylamide Gel Preparation Using N-hydroxysuccinimide-acrylamide Ester to Study Cell-Extracellular Matrix Mechanical Interactions. Frontiers in Materials, 8, February 2021.

27. Prabu Karthick Parameshwar, Lucas Sagrillo-Fagundes, Caroline Fournier, Sylvie Girard, Cathy Vaillancourt, and Christopher Moraes. Disease-specific extracellular matrix composition regulates placental trophoblast fusion efficiency. Biomaterials Science, 9(21):7247– 7256, October 2021.

28. Seiichiro Ishihara, Haruna Kurosawa, and Hisashi Haga. Stiffness-Modulation of Collagen Gels by Genipin-Crosslinking for Cell Culture. Gels, 9(2):148, February 2023.

29. Sana Syed, Amin Karadaghy, and Silviya Zustiak. Simple Polyacrylamide-based Multiwell Stiffness Assay for the Study of Stiffness-dependent Cell Responses. Journal of Visualized Experiments: JoVE, (97):52643, March 2015.

30. Yeling Ma, Qian Yang, Mengjie Fan, Lanmei Zhang, Yan Gu, Wentong Jia, Zhilang Li, Feiyang Wang, Yu-xia Li, Jian Wang, Rong Li, Xuan Shao, and Yan-Ling Wang. Placental endovascular extravillous trophoblasts (enEVTs) educate maternal T-cell differentiation along the maternal-placental circulation. Cell Proliferation, pages e12802–e12802, April 2020.

31. Amnon Buxboim, Karthikan Rajagopal, Andre’ E.X. Brown, and Dennis E. Discher. How deeply cells feel: methods for thin gels. Journal of physics. Condensed matter: an Institute of Physics journal, 22(19):194116, May 2010.

32. Paul A. Janmey, Daniel A. Fletcher, and Cynthia A. Reinhart-King. Stiffness Sensing by Cells. Physiological Reviews, 100(2):695–724, April 2020.

33. Aleksandra K. Denisin and Beth L. Pruitt. Tuning the Range of Polyacrylamide Gel Stiffness for Mechanobiology Applications. ACS Applied Materials & Interfaces, 8(34):21893–21902, August 2016.

34. Vonetta L Edwards, Elias McComb, Jason P Gleghorn, Larry Forney, Patrik M Bavoil, and Jacques Ravel. Three-dimensional models of the cervicovaginal epithelia to study host–microbiome interactions and sexually transmitted infections. Pathogens and Disease, 80(1):ftac026, August 2022.

35. N’Dea S. Irvin-Choy, Katherine M. Nelson, Jason P. Gleghorn, and Emily S. Day. Design of nanomaterials for applications in maternal/fetal medicine. Journal of Materials Chemistry B, May 2020.

36. Katherine M. Nelson, N’Dea Irvin-Choy, Matthew K. Hoffman, Jason P. Gleghorn, and Emily S. Day. Diseases and conditions that impact maternal and fetal health and the potential for nanomedicine therapies. Advanced Drug Delivery Reviews, 170:425–438, March 2021.

